# The Brain Interactome of a Permissive Prion Replication Substrate

**DOI:** 10.1101/2024.08.19.608704

**Authors:** Hamza Arshad, Shehab Eid, Surabhi Mehra, Declan Williams, Lech Kaczmarczyk, Erica Stuart, Walker S. Jackson, Gerold Schmitt-Ulms, Joel C. Watts

**Author notes:** To whom correspondence should be addressed at: Krembil Discovery Tower, Rm. 4KD481, 60 Leonard Ave., Toronto, ON, Canada, M5T 0S8.

## Abstract

Bank voles are susceptible to prion strains from many different species, yet the molecular mechanisms underlying the ability of bank vole prion protein (BVPrP) to function as a universal prion acceptor remain unclear. Potential differences in molecular environments and protein interaction networks on the cell surface of brain cells may contribute to BVPrP’s unusual behavior. To test this hypothesis, we generated knock-in mice that express physiological levels of BVPrP (M109 isoform) and employed mass spectrometry to compare the interactomes of mouse (Mo) PrP and BVPrP following mild *in vivo* crosslinking of brain tissue. Substantial overlap was observed between the top interactors for BVPrP and MoPrP, with established PrP-interactors such as neural cell adhesion molecules, subunits of Na^+^/K^+^-ATPases, and contactin-1 being equally present in the two interactomes. We conclude that the molecular environments of BVPrP and MoPrP in the brains of mice are very similar. This suggests that the unorthodox properties of BVPrP are unlikely to be mediated by differential interactions with other proteins.

## Introduction

Prion diseases such as scrapie in sheep, bovine spongiform encephalopathy, chronic wasting disease (CWD) in cervids, and Creutzfeldt-Jakob disease (CJD) in humans are caused by the accumulation of misfolded prion protein (PrP) in the brain. Encoded by the *Prnp* gene in rodents, PrP is a glycosylphosphatidylinositol (GPI)-anchored protein present on the cell surface of neurons and astrocytes that is modified post-translationally by the addition of up to two N-linked glycans. During prion disease, PrP undergoes conversion from its normal, cellular form (PrP^C^) into a self-propagating, aggregation-prone conformer termed PrP^Sc^ (1). These two PrP conformers have distinct physicochemical properties: PrP^C^ is a predominantly α-helical and sensitive to digestion with proteases whereas PrP^Sc^ is composed almost entirely of β-sheets and, due to its aggregated nature, is partially resistant to digestion with proteases such as proteinase K (2–5). Prion diseases are transmissible, as an exogenous source of PrP^Sc^ can template the conversion of host expressed PrP^C^ into additional copies of PrP^Sc^, allowing prions to spread both within and to the brain via a cascade of protein misfolding. PrP^C^ expression is necessary for both prion replication and prion neurotoxicity (6–9). Prion disease pathogenesis likely stems directly from PrP^Sc^ accumulation or prion replication rather than loss of PrP^C^ since PrP knockout (PrP^-/-^) mice do not succumb to prion disease during their lifespans (10).

While intraspecies prion transmission is generally efficient, cross-species transmission is typically an inefficient process characterized by incomplete disease transmission and prolonged incubation periods. This phenomenon, known as the “species barrier”, is mediated by both amino acid sequence differences between PrP^Sc^ and PrP^C^ as well as structural compatibility between the two molecules (11–13). Bank voles (*Myodes glareolus*) are atypical in that they do not impose a substantial species barrier during prion transmission and are highly susceptible to prions from many different species (14–21). Transgenic or knock-in mice expressing bank vole PrP (BVPrP) recapitulate the enhanced prion susceptibility of bank voles, indicating that this phenomenon is mediated by BVPrP itself (22–24). Moreover, BVPrP functions as a highly permissive prion substrate when expressed in cultured cells or used for *in vitro* prion conversion assays (25–31). Collectively, these results suggest that BVPrP may be capable of functioning as a “universal acceptor” for prions.

A molecular explanation for the enhanced ability of BVPrP to mediate cross-species prion transmission remains to be fully determined. The sequence of mature BVPrP, following removal of N- and C-terminal signal sequences, differs from that of mouse PrP (MoPrP) at only eight positions, suggesting that a small number of residues can have a large effect on the properties of the protein. Indeed, the ability of BVPrP to enable replication of both mouse and hamster prion strains is governed by five key residues (32). Additionally, BVPrP is polymorphic at codon 109, where either a methionine (M109) or isoleucine (I109) residue can be present (33). Transgenic mice over-expressing wild-type (WT) or mutant BVPrP(I109) as well as knock-in mice expressing physiological levels of mutant BVPrP(I109) develop spontaneous disease that exhibits many of the hallmarks of authentic prion disease, suggesting that BVPrP may be intrinsically prone to adopting misfolded conformations (34–37).

The issue of whether proteins other than PrP participate in prion replication *in vivo* remains an open question (38). Non-protein co-factors such as polyanions and specific lipids can modulate prion replication and the properties of prion strains *in vitro* (39–43). Based on prion transmission experiments, it has been hypothesized that a PrP-interacting protein, provisionally designated “Protein X”, may be required for prion replication (44, 45). Indeed, PrP^C^ is known to interact with a number of other cell membrane proteins including neural cell adhesion molecule (NCAM1 and NCAM2) (46, 47), subunits of Na^+^/K^+^-ATPases (NKAs) (48, 49), dipeptidyl aminopeptidase-like protein 6 (DPP6) (50), the laminin receptor (51), and G protein-couple receptor 126 (Gpr126, also known as Adgrg6) (52). Interactome experiments have revealed many more proteins residing in close spatial proximity to PrP^C^ within the membrane (48, 53–57). Although identified PrP^C^-interacting proteins have yet to provide insight into the conversion of PrP^C^ into PrP^Sc^, they have revealed that PrP^C^ may influence multiple biological processes within the cell (58, 59).

We hypothesized that due to its unique amino acid composition, BVPrP may interact with a distinct set of proteins in the brain compared to MoPrP, and that this may in part explain the ability of BVPrP to promote cross-species prion transmission. To address this issue experimentally, we compared the interactomes of MoPrP and BVPrP using healthy WT C57BL/6 mice expressing MoPrP and knock-in mice expressing BVPrP(M109). Qualitative and quantitative mass spectrometry experiments revealed that the molecular environments of MoPrP and BVPrP within the brain are very similar, arguing that intrinsic biophysical features of BVPrP rather than altered protein-protein interactions drive the ability of BVPrP to function as a universal prion acceptor.

## Results

Using an identical strategy to what we recently used to generate knock-in (ki) mice expressing WT or mutant BVPrP(I109) (37), we generated ki mice expressing WT BVPrP(M109), referred to hereafter as kiBVM mice. To create kiBVM mice, the PrP open reading frame within Exon 3 of mouse *Prnp* was replaced with the corresponding region of bank vole *Prnp*, allowing BVPrP expression to be controlled by the endogenous mouse *Prnp* promoter (**Fig. 1A**). PrP levels in brain homogenates from kiBVM, WT C57BL/6 mice, and co-isogenic PrP^-/-^ mice on a C57BL/6 background (60) were compared by immunoblotting. Using three different antibodies (SAF32, HuM-D13, and POM1) that recognize different epitopes within PrP (**Fig. 1A**), the expression level of BVPrP in the brains of kiBVM mice and MoPrP in the brains of WT mice were found to be similar **(Fig. 1B)**. As expected, the HuM-R1 antibody, which recognizes an epitope uniquely present in the C-terminal region of MoPrP, failed to detect BVPrP in the brains of kiBVM mice. None of the antibodies detected PrP signal in brain homogenates from PrP^-/-^ mice. Under physiological conditions, PrP^C^ undergoes endoproteolytic cleavage in the vicinity of residues 110/111 to release an N-terminal N1 fragment, leaving behind a membrane-anchored C-terminal fragment termed C1 (61). Following the removal of N-linked glycans from PrP using PNGase F, the C1 fragment was observed in brain homogenates from both WT and kiBVM mice (**Fig. 1C**). Brain BVPrP levels did not change with age in kiBVM mice, and there were no apparent differences in brain BVPrP expression levels between male and female mice (**Fig. 1D**).

**Figure 1.**
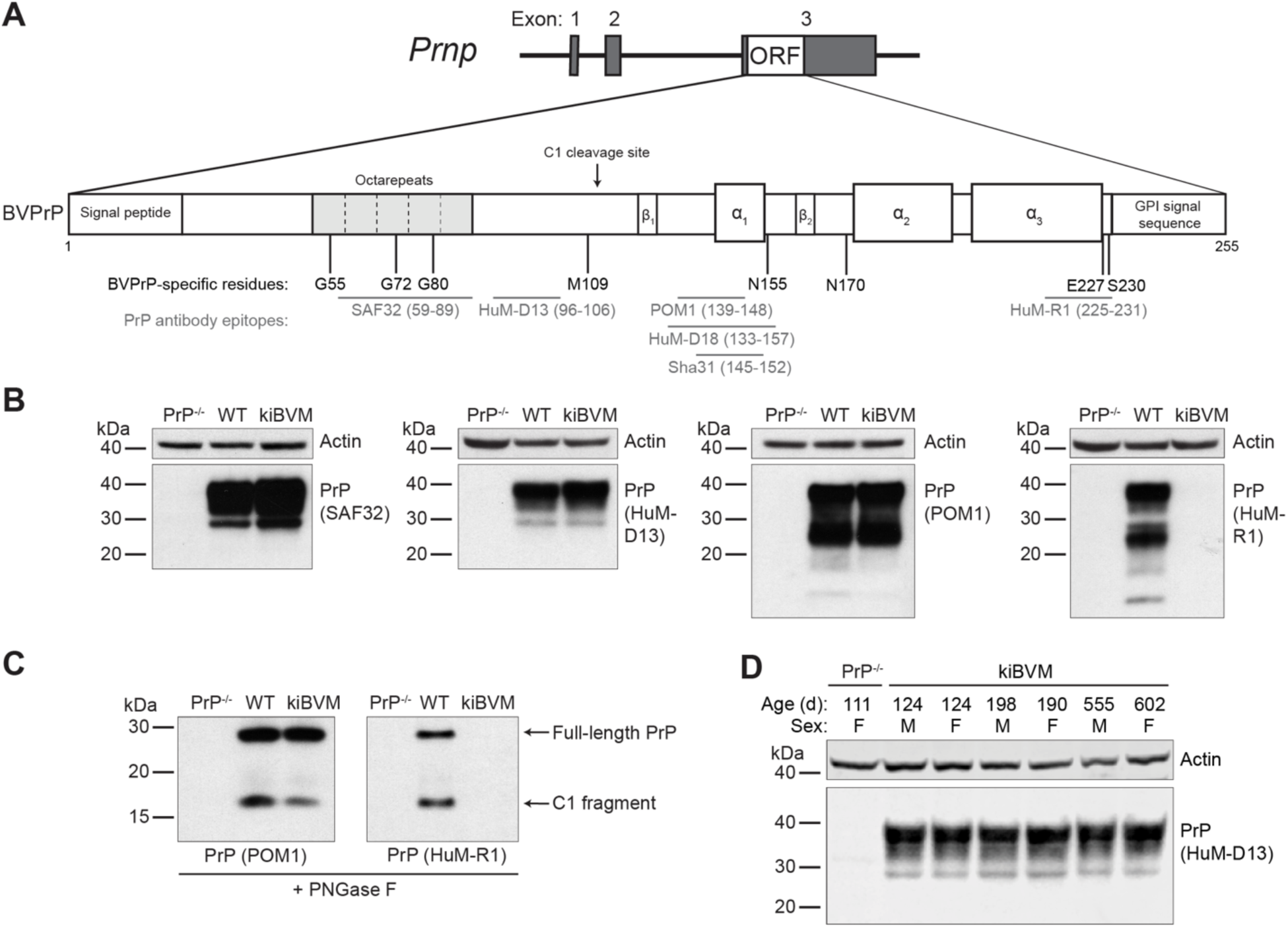
Generation of knock-in mice expressing bank vole PrP. **A**) Schematic of the gene-targeted allele in kiBVM mice expressing BVPrP(M109). The eight amino acid residue differences between the mature forms of BVPrP and MoPrP are shown, as are the approximate epitopes for the anti-PrP antibodies used in this study. **B**) Immunoblots for PrP in brain extracts from healthy PrP^-/-^, wild-type C57BL/6 (WT), and kiBVM mice probed with antibodies that recognize both MoPrP and BVPrP (SAF32, HuM-D13, and POM1) or only MoPrP (HuM-R1). Blots were reprobed with an antibody against actin. **C**) Immunoblots for PrP in PNGase F-treated brain extracts from PrP^-/-^, WT, and kiBVM mice probed with the antibodies POM1 or HuM-R1. Full-length BVPrP as well as the C1 endoproteolytic product are indicated. **D**) Immunoblot for PrP in brain extracts from kiBVM mice at the indicated ages probed with the antibody HuM-D13. Brains from both male (M) and female (F) mice were analyzed. The blot was reprobed with an antibody against actin.

We also examined PrP expression levels in peripheral tissues from WT and kiBVM mice. Using an antibody that does not detect the C1 fragment, comparable levels of PrP expression were found in brain, spinal cord, stomach, spleen, skin, testis, lung, heart, muscle, and tongue tissue between the two mouse lines (**Fig. 2A**). Varying amounts of diglycosylated, monoglycosylated, and unglycosylated PrP species were observed across the different tissues. Interestingly, an increase in PrP species that migrated more rapidly was found when analyzing testis tissue. This likely corresponds to the C2 endoproteolytic fragment, which results from cleavage of PrP near residue 90. Using an antibody that detects full-length, C1, and C2 PrP species, PrP expression levels in peripheral tissues were found to be much lower than those observed in the brain and spinal cord **(Fig. 2B)**. Endoproteolytic processing of PrP to produce the C1 fragment was observed in brain, spinal cord, heart, muscle, tongue, and skin tissue, but not in the lungs or testis. Overall, we conclude that kiBVM mice express physiological levels of PrP, both in the central nervous system and in peripheral tissues.

**Figure 2.**
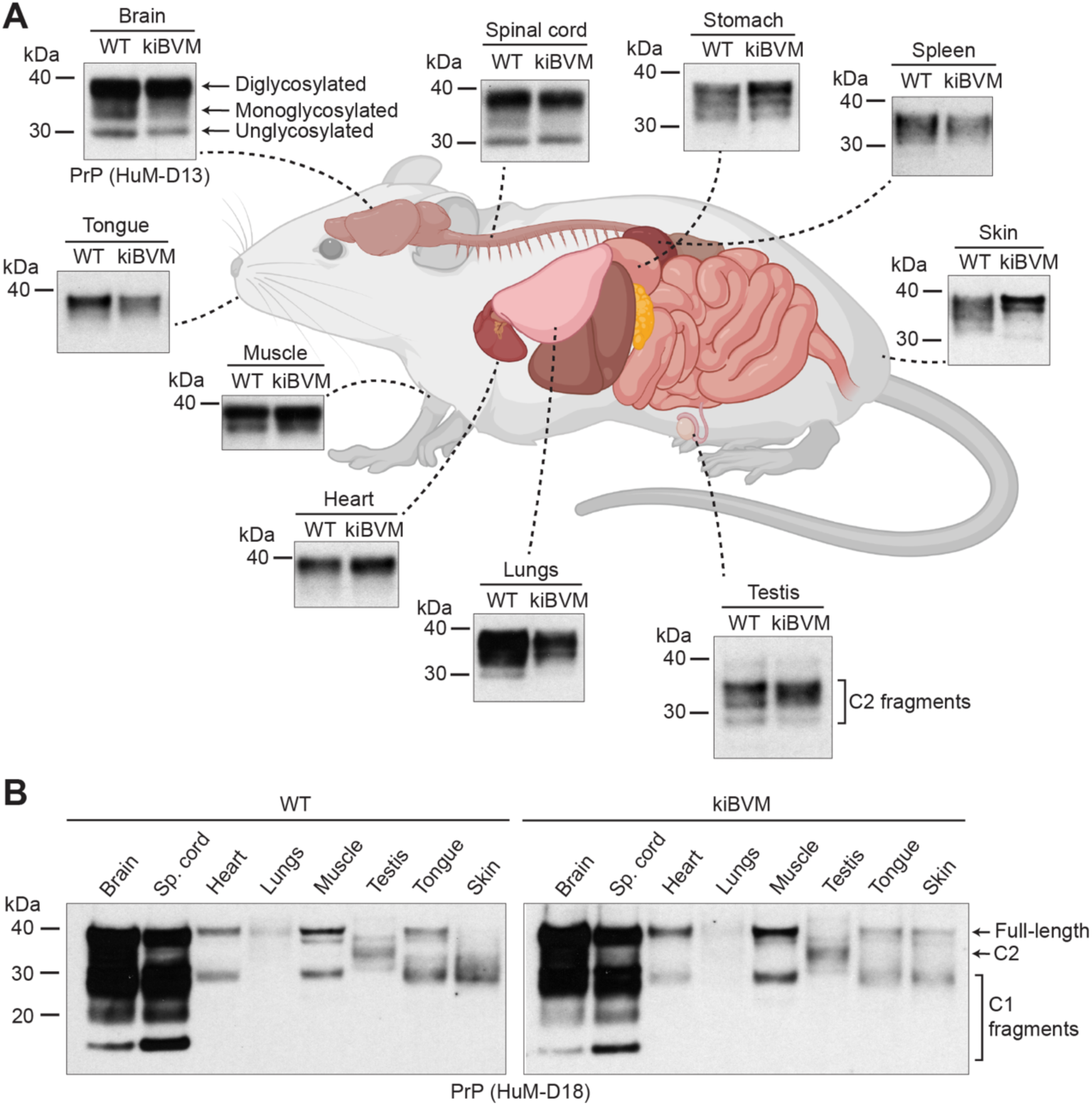
Tissue expression profile of PrP in wild-type and BVPrP knock-in mice. **A**) Immunoblots for PrP in extracts prepared from brain, spinal cord, stomach, spleen, skin, testis, lung, heart, tongue, and muscle tissue from healthy WT C57BL/6 and kiBVM mice. For the brain tissue immunoblot, di-, mono-, and unglycosylated PrP species are indicated. For the testis immunoblot, putative C2 endoproteolytic products are marked. PrP was detected using the antibody HuM-D13. The mouse organ schematic was generated using BioRender.com. **B**) Immunoblots for relative PrP expression levels in the indicated tissues from either WT C57BL/6 (left blot) or kiBVM (right blot) mice. Full-length PrP as well as the C1 and C2 endoproteolytic fragments are indicated. PrP was detected using the antibody HuM-D18.

Transgenic mice overexpressing WT BVPrP(I109) spontaneously develop prion disease as they age (34, 35). In contrast, knock-in mice expressing physiological levels of WT BVPrP(I109) remain healthy up to 20 months of age (37). One study reported that transgenic mice overexpressing BVPrP(M109) develop a spontaneous but non-transmissible proteinopathy that lacks biochemical features associated with prion disease such as the presence of protease-resistant PrP in the brain (24). Thus, we asked whether aged kiBVM mice develop spontaneous disease. Cohorts of 9-10 kiBVM and WT mice were monitored longitudinally for the development of clinical signs of neurological illness. All kiBVM and WT mice remained free of neurological illness up to 20 months of age (**Fig. 3A**). As expected, PrP levels in the brain did not differ between WT and kiBVM mice at 20 months of age (**Fig. 3B**). On average, levels of detergent-insoluble PrP species in brain homogenates from 20-month-old kiBVM mice were approximately 25% lower than in age-matched brain homogenates from WT mice (**Fig. 3C, D**). As a positive control, we included brain homogenates from spontaneously ill knock-in mice expressing D178N- or E200K-mutant BVPrP(I109) (37). Despite lower levels of total PrP expression (**Fig. 3B**), increased amounts of detergent-insoluble PrP were present in the brains of spontaneously sick mice expressing mutant BVPrP(I109) compared to asymptomatic 20-month-old WT and kiBVM mice (**Fig. 3C**). Finally, we looked for the presence of thermolysin (TL)-resistant PrP species in brain homogenates from aged WT and kiBVM mice. Consistent with previous findings (37), TL-resistant PrP could be detected in brain homogenates from spontaneously ill knock-in mice expressing mutant BVPrP(I109) (**Fig. 3E**). However, no detergent-insoluble TL-resistant PrP species were observed in the brains of any of the aged WT or kiBVM mice. We therefore conclude that kiBVM mice do not develop spontaneous disease or exhibit misfolded PrP species in their brains with aging.

**Figure 3.**
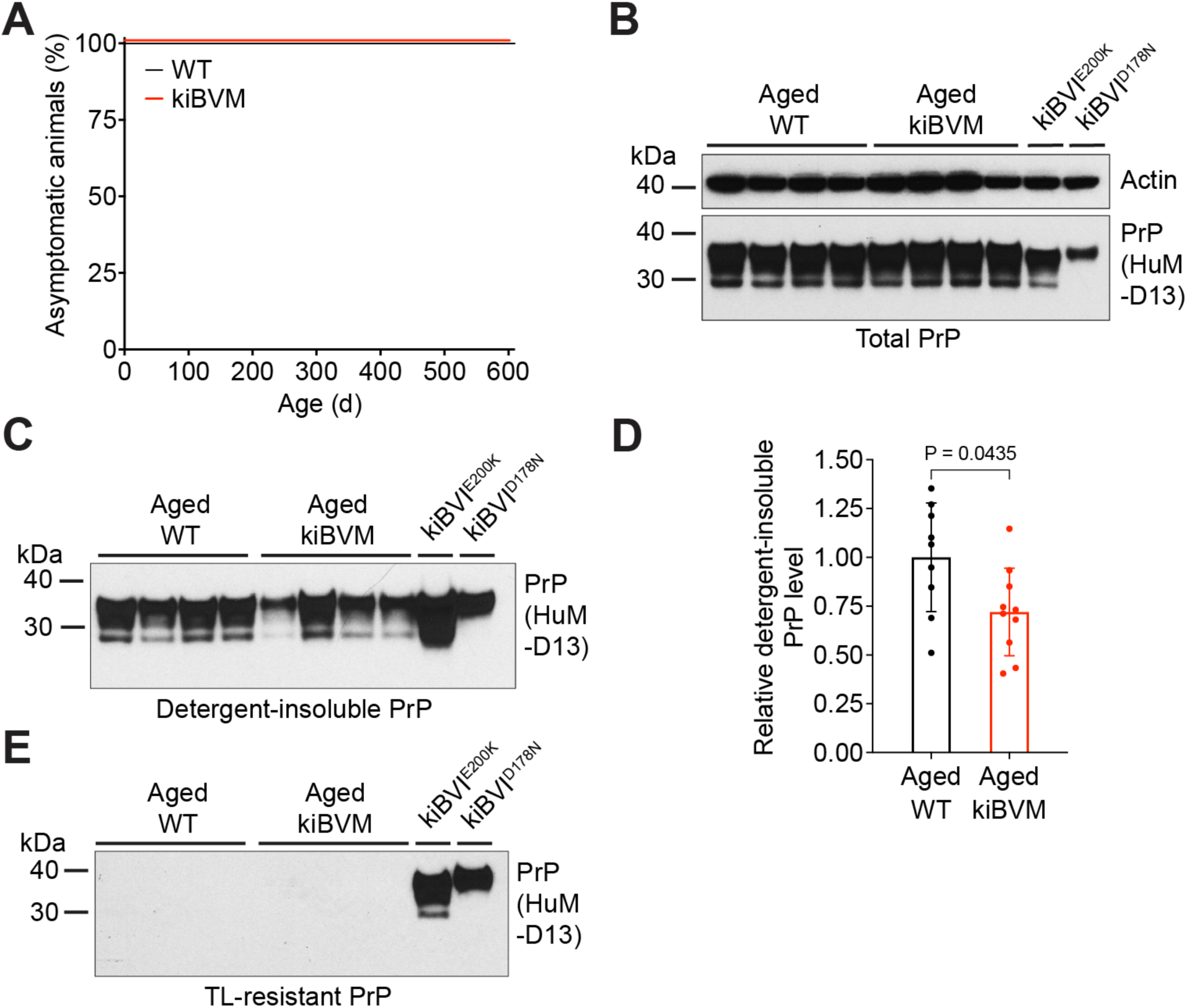
Knock-in mice expressing BVPrP(M109) do not develop spontaneous disease. **A**) Kaplan-Meier survival curves for WT C57BL/6 (black line; n = 9) and kiBVM (red line; n = 10) mice. **B**) Immunoblot of total PrP levels in detergent-extracted brain homogenates from aged asymptomatic (18-to 20-months-old) WT C57BL/6 and kiBVM mice (n = 4 each). Brain homogenates from spontaneously ill kiBVI^E200K^ and kiBVI^D178N^ mice are shown as controls. The blot was reprobed with an antibody against actin. **C**) Immunoblot of detergent-insoluble PrP levels in brain homogenates from aged asymptomatic WT C57BL/6 and kiBVM mice (n = 4 each). Brain homogenates from spontaneously ill kiBVI^E200K^ and kiBVI^D178N^ mice are shown as controls. **D**) Quantification of detergent-insoluble PrP levels in brain homogenates from aged asymptomatic WT C57BL/6 (n = 9) and kiBVM (n = 10) mice. Statistical significance was assessed using a Mann-Whitney test. **E**) Immunoblot for thermolysin (TL)-resistant PrP in brain homogenates from aged asymptomatic WT C57BL/6 and kiBVM mice (n = 4 each). Brain homogenates from spontaneously ill kiBVI^E200K^ and kiBVI^D178N^ mice are shown as controls. In panels B, C, and E, PrP was detected using the antibody HuM-D13.

The interactome of a protein can be considered the complete set of proteins with which that protein interacts. Previously, the interactome of mouse PrP^C^ in brain tissue and cultured cells has been investigated using time-controlled transcardiac perfusion crosslinking (tcTPC) (48, 53, 54, 57). In this technique, proteins residing in close spatial proximity are covalently linked together by perfusing mice with formaldehyde prior to brain removal. Immunoprecipitation is used to isolate PrP^C^-containing crosslinked protein complexes and then, after reduction, alkylation, and trypsinization, samples are subjected to liquid chromatography followed by tandem mass spectrometry (LC-MS/MS) to identify potential PrP^C^-interacting proteins (**Fig. 4A**). To compare the interactomes of BVPrP and MoPrP, 3 kiBVM, 3 WT, and 2 PrP^-/-^ mice at 3 months of age were subjected to tcTPC, their brains were removed and homogenized, and then PrP^C^-containing complexes were isolated by immunoprecipitation using the HuM-D18 antibody. An immunoblot confirmed the successful immunoprecipitation of crosslinked brain PrP^C^-containing complexes, which are indicated by the presence of high molecular weight smears in the eluates from WT and kiBVM, but not PrP^-/-^ mice (**Fig. 4B**).

**Figure 4.**
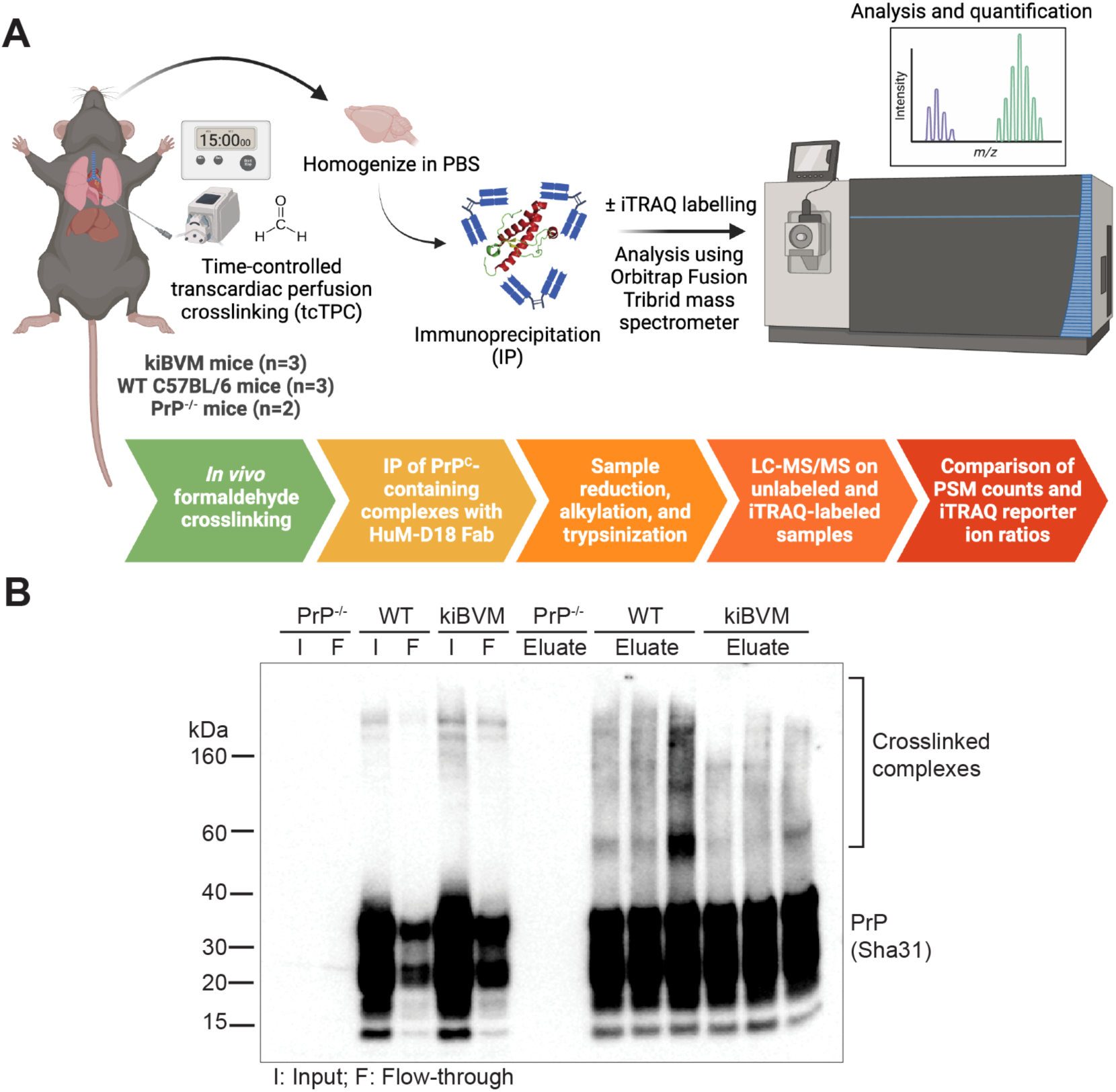
Workflow for interactome experiments following time-controlled transcardiac perfusion crosslinking in mice. **A**) Schematic workflow for the mass spectrometry-based comparative PrP^C^ interactome experiments in the brains of kiBVM, WT C57BL/6, and PrP^-/-^ mice. The schematic was generated using BioRender.com. **B**) Immunoblot for PrP in brain homogenates from PrP^-/-^, WT C57BL/6, and kiBVM mice subjected to tcTPC and then immunoprecipitated using the anti-PrP antibody HuM-D18. The fractions shown are the input (I; pre-immunoprecipitation), flow-through (F; non-bound fraction following immunoprecipitation), and eluate (bound fraction following immunoprecipitation). PrP was detected with Sha31 antibody.

Following LC-MS/MS, potential PrP^C^-interacting proteins were identified by applying stringent criteria that required each protein to have been identified by at least 2 unique peptide-to-spectrum matches (PSMs) and that the ratio of the number of PSMs identified in the kiBVM samples to the number of PSMs identified in the PrP^-/-^ samples was greater than 3. Hits were then ranked based on the average number of PSMs in the kiBVM samples minus the average number of PSMs in the PrP^-/-^ samples. The top 30 identified proteins from the brains of kiBVM mice are shown in **Fig. 5**. Not surprisingly, the top hit in the BVPrP interactome was the prion protein itself, indicating that the targeted pull-down of PrP^C^-containing complexes was successful. Other confidently identified hits included NCAM1 and NCAM2; the α1, α2, α3, and β1 subunits of NKAs; DPP6; contactin-1; the protein disulfide isomerase P4HB; and synaptic vesicle glycoprotein 2B (SV2B); all of which have been identified in previous PrP^C^ interactome studies (48, 53, 54, 57). For each of the the top 30 proteins identified in the BVPrP interactome, we also computed the average number of PSMs identified in samples from WT mice minus the average number of PSMs from the PrP^-/-^ mice. This analysis revealed that 27 of 30 proteins were also present in the MoPrP interactome dataset, indicative of substantial overlap between the BVPrP and MoPrP interactomes (**Fig. 5**). The remaining 3 proteins were also identified but didn’t reach the specificity threshold of a WT:PrP^-/-^ PSM ratio of at least 3. We also computed the MoPrP interactome by ranking the identified proteins based on their enrichment in WT mice (**Fig. S1**). Remarkably, 25 of the top 30 proteins identified in the MoPrP interactome were also top 30 hits in the BVPrP interactome, and the other 5 proteins were present in the BVPrP interactome but did not make the top 30.

**Figure 5.**
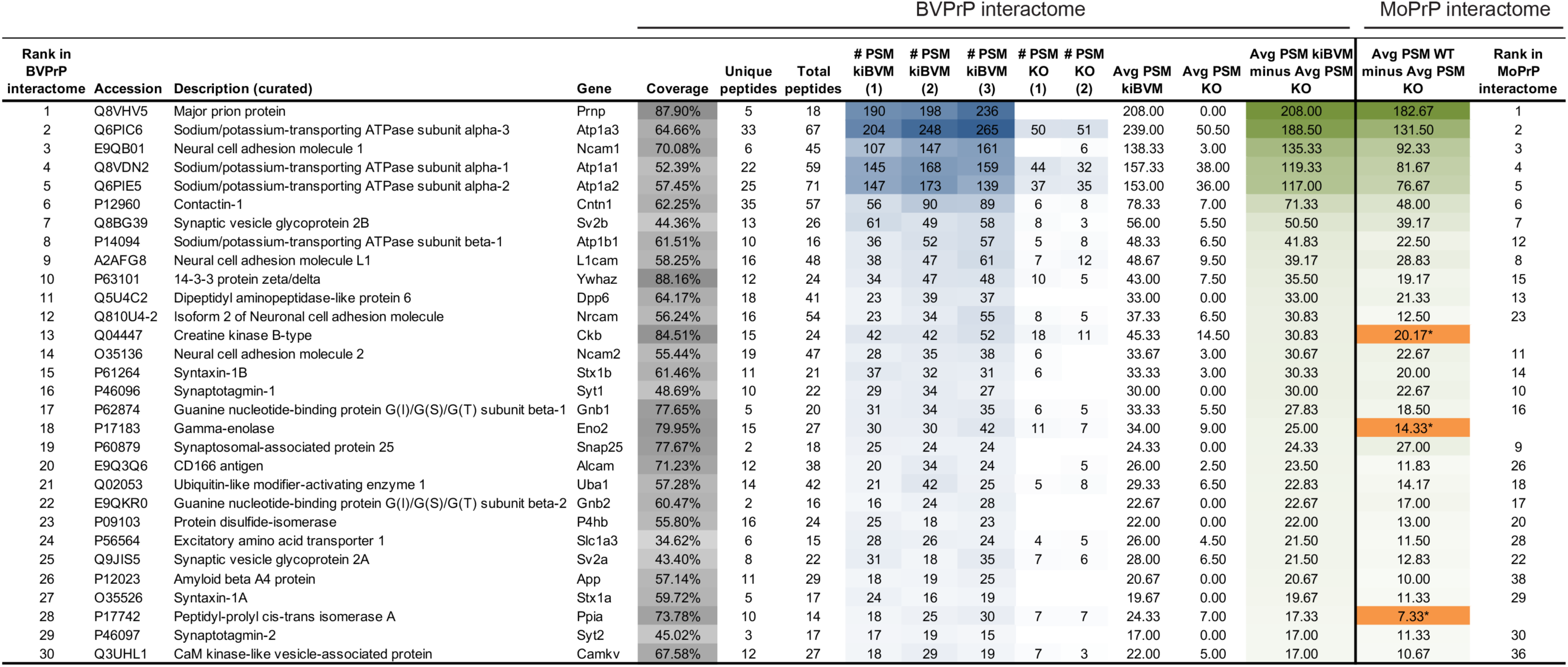
The brain interactome of bank vole PrP. Top 30 identified BVPrP-interacting proteins ranked by the average number of peptide-to-spectrum matches (PSMs) observed in kiBVM mice minus the average number of PSMs observed in PrP^-/-^ mice (KO). For each protein identified, the average number of PSMs observed in WT mice minus the average number of PSMs from PrP^-/-^ mice as well as the rank in the MoPrP interactome are also shown. Coverage, represented by grey shading, indicates the percentage of primary structure covered by PSMs across the three kiBVM samples. Blue-shaded numbers reflect the number of PSMs detected and green-shaded numbers reflect the relative enrichment in kiBVM or WT versus PrP^-/-^ samples. Orange shading with an asterisk indicates those proteins that were also found in the MoPrP interactome but didn’t reach the specificity threshold (WT:KO PSM ratio of at least 3).

To determine whether any of the identified proteins may associate more strongly with either BVPrP or MoPrP, we performed quantitative LC-MS/MS analysis on the tcTPC-treated brain samples using isobaric tags for relative and absolute quantitation (iTRAQ) (62). In the previous analyses using unlabeled samples, quantitative assessment of individual protein hits was not possible since the mass spectrometry runs on the immunoprecipitated brain samples were conducted independently. Following reduction, alkylation, and trypsin digestion, each of the 8 pull-down samples was labelled with a different 8-plex iTRAQ reagent and then combined and analyzed together by LC-MS/MS. iTRAQ reporter ion ratios relative to one of the two PrP^-/-^ samples were calculated to determine the relative enrichment of specific peptides in the different samples. Peptides corresponding to PrP were strongly enriched in all the pull-down samples from the brains of both kiBVM and WT mice, but not in the sample from the other PrP^-/-^ brain (**Fig. 6A**, top panel). When only peptides containing at least one of the eight BVPrP-specific residues (i.e., not present in the MoPrP sequence) were considered, iTRAQ reporter ion enrichment was only observed for the samples from kiBVM mice (**Fig. 6A**, lower panel). Conversely, when only peptides containing at least one MoPrP-specific residue (i.e., not present in the BVPrP sequence) were analyzed, enrichment was only observed in the samples from WT mice (**Fig. S2**). A non-specific interactor such as glyceraldehyde-3-phosphate dehydrogenase (GAPDH), which likely associates with the immunoprecipitation matrix, did not exhibit any consistent iTRAQ reporter ion enrichment across the kiBVM, WT, and PrP^-/-^ samples (**Fig. S3**). iTRAQ reporter ion ratios for the top putative PrP^C^-interacting proteins were compared between the kiBVM and WT samples. Although there was considerable variability between samples, there were no significant differences in the relative enrichments between kiBVM and WT mice for any of the proteins (**Fig. 6B**). Thus, the quantitative iTRAQ labelling data is consistent with the unlabeled experiments where PSM counts were compared, with both sets of experiments suggesting that the interactomes of BVPrP and MoPrP in healthy mice are highly congruent.

**Figure 6.**
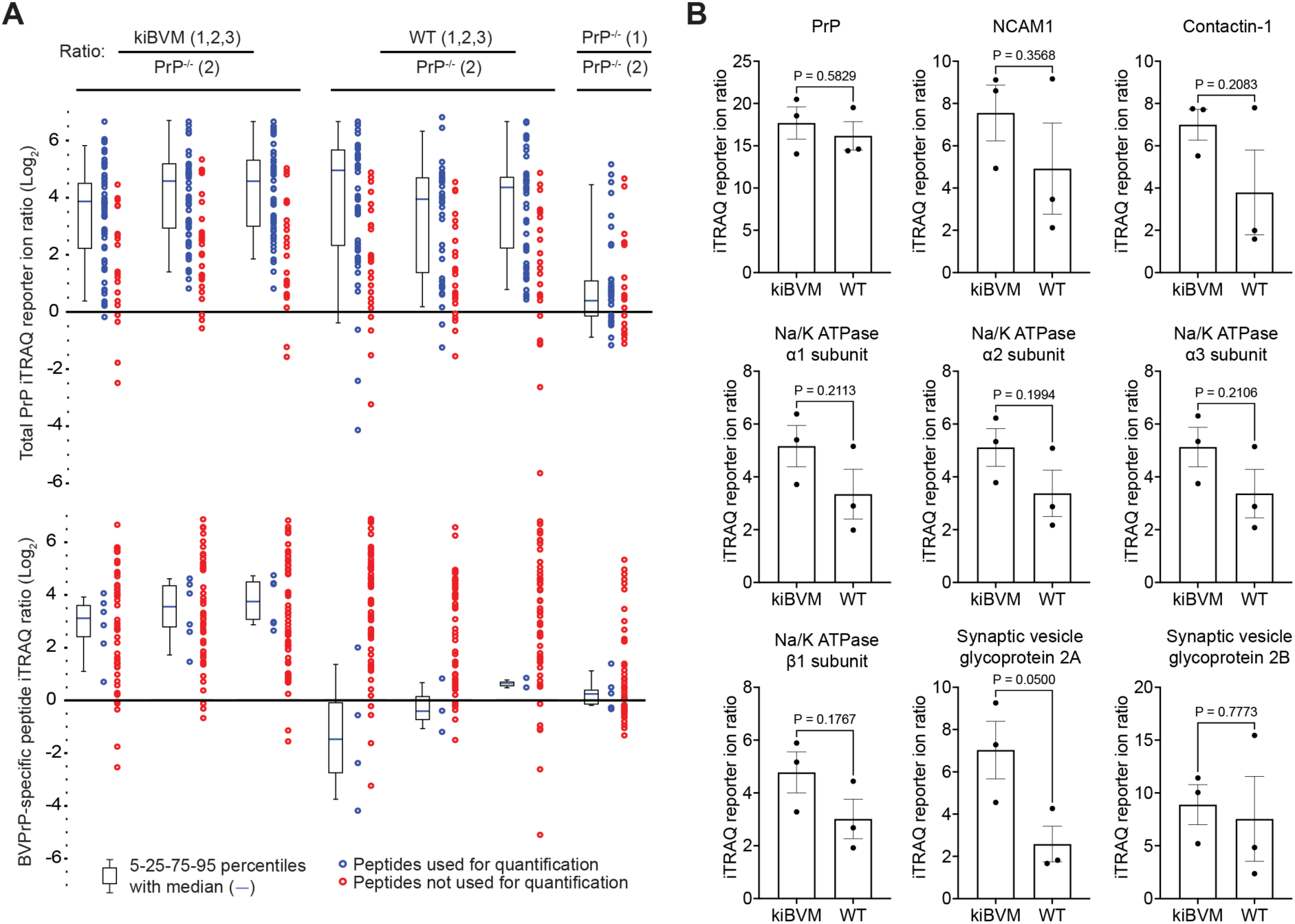
Quantitative analysis of mouse and bank vole PrP interactors using iTRAQ labelling. **A**) Boxplots depicting iTRAQ reporter ion enrichment ratios (Log_2_) for PrP peptides in immunoprecipitated samples from the brains of kiBVM and WT C57BL/6 mice (n = 3 each) as well as the brain of a PrP^-/-^ mouse. For each sample, the enrichment ratio was calculated in comparison to the second PrP^-/-^ sample. The upper graph displays enrichment ratios for all PrP peptides, whereas the bottom panel displays enrichment ratios for PrP peptides containing at least one of the eight BVPrP-specific amino acids. Peptides that were not used or quantification include those that did not pass the Percolator threshold and those that were excluded to prevent double counting when assigned correctly by both the Sequest and Mascot algorithms in Proteome Discoverer. **B**) iTRAQ reporter ion ratios (Log_2_), all in comparison to the second PrP^-/-^ sample, for a subset of proteins identified in the kiBVM and WT interactome datasets (n = 3 each). Statistical significance was assessed using two-tailed unpaired *t*-tests.

## Discussion

Transgenic mice expressing BVPrP retain the ability of bank voles to function as a universal prion acceptor, suggesting that the primary sequence of BVPrP is the main driver of its prion permissive behavior (22, 23). Additionally, this suggests that if altered protein-protein interactions, mediated by BVPrP-specific residues, are responsible for facilitating cross-species prion replication, these interactions must be conserved in mice. In this study, we compared the interactomes of MoPrP and BVPrP in mice to discern any novel protein-protein interactions that could explain the heightened susceptibility of bank voles to prions. To avoid potential artifacts associated with PrP^C^ overexpression in transgenic mice, we generated knock-in mice that express BVPrP(M109) at physiological levels under the control of the endogenous mouse *Prnp* promoter. This ensured that MoPrP and BVPrP were expressed at the same levels in the brains of mice and with the same spatiotemporal pattern of expression. The interactomes of MoPrP and BVPrP were remarkably similar, with no unique interactors identified for either protein. This suggests that the molecular environments of MoPrP and BVPrP within the plasma membrane of brain cells are highly similar, if not identical.

Although it has been firmly established that BVPrP(I109) is prone to spontaneous misfolding (34–37, 42, 63), it is less clear whether BVPrP(M109) can also form prions spontaneously. In one study, transgenic mice overexpressing BVPrP(M109) developed clinical disease, but without accompanying biochemical features characteristic of prion disease, such as the presence of protease-resistant PrP (24). No disease transmission was observed upon inoculation of brain extracts from spontaneously sick BVPrP(M109) transgenic mice into knock-in mice expressing BVPrP(M109), suggesting that overexpression of BVPrP(M109) causes a PrP proteinopathy without spontaneous prion formation. We found that, like WT mice expressing MoPrP, kiBVM mice remained healthy throughout their normal lifespans without any evidence for an increase in misfolded BVPrP species in the brains of aged mice. This is identical to what we observed in knock-in mice expressing the more misfolding-prone BVPrP(I109) (37), arguing that BVPrP overexpression is required to elicit spontaneous disease and prion generation, which is consistent with absence of reported spontaneous prion disease in bank voles.

The issue of whether proteins other than PrP participate in prion formation or replication remains an unsolved mystery (38). Given its prion permissive nature, we speculated that BVPrP may represent an ideal means for identifying such proteins. However, both our unlabeled and iTRAQ-labeled interactome datasets failed to reveal proteins that selectively interact with BVPrP. Moreover, since we also did not detect any proteins that selectively interact with MoPrP, this makes a scenario where the enhanced prion replication properties of BVPrP stem from a reduced ability to interact with a negative regulator of prion replication less plausible. Nonetheless, our interactome studies did reveal that BVPrP interacts with many previously identified MoPrP interactors. A top hit for both BVPrP and MoPrP was NCAM1, which has been robustly identified as a PrP^C^-interacting protein in several lines of cultured cells and in mice (46, 53, 54, 57). NCAM1 is unlikely to be involved in prion replication since NCAM1 knockout mice are as susceptible to prions as WT mice (46). In contrast, the interaction between NCAM1 and PrP^C^, mediated by the flexible N-terminal domain of PrP^C^ and the fibronectin type-3 domain of NCAM, has been shown to promote neurite outgrowth and neuronal differentiation (47, 64, 65). PrP^C^ also controls the polysialylation of NCAM1 in cultured cells, a phenomenon that is important for epithelial-to-mesenchymal transition during cellular development (66). Another robust hit for both BVPrP and MoPrP were subunits of NKAs, the pumps that are responsible for maintaining the resting potential across the plasma membrane in neural cells. PrP^C^ co-localizes with the NKA α1 subunit in neural cells and promotes NKA-mediated ion uptake activity in neuronal and astroglial cultures (48, 49). Subunits of NKAs have also been found in preparations of scrapie-associated fibrils isolated from brains of prion-infected mice (67). Interestingly, targeting the NKA α1 subunit with cardiac glycosides results in their internalization as well as the co-internalization and degradation of PrP^C^ (68). Thus, reducing PrP^C^ levels by targeting the NKA α subunit is a potential strategy for dampening prion replication during prion disease (69).

The complexity of the lipid bilayer within which PrP^C^ resides complicates the identification of PrP^C^ binding partners. Our experiments, which relied on the mild crosslinking of PrP^C^ to its nearest neighbors, revealed proteins that are found in the vicinity of PrP^C^, and parallel analyses in PrP^-/-^ mice were useful for delineating specific from non-specific interactions. However, because crosslinking was used, it is possible that not all identified proteins physically interact with PrP^C^ and may simply reside in close spatial proximity within the membrane. Also, our study does not discriminate between direct and indirect interactors of PrP^C^, and it is likely that the crosslinking followed by immunoprecipitation approach may not have captured all biologically meaningful protein-protein interactions involving PrP^C^. Another limitation of our study is that we only investigated the PrP^C^ interactome in healthy mice. It remains possible that BVPrP may participate in a unique interaction with another protein that only occurs during the conversion of PrP^C^ into PrP^Sc^, thus facilitating cross-species prion infection. It is also conceivable that BVPrP may exhibit enhanced binding to a non-proteinaceous cofactor that may assist with the conversion of PrP^C^ into PrP^Sc^. Indeed, lipid cofactors such as phosphatidylethanolamine have been shown to enhance the templated or spontaneous conversion of mouse PrP^C^ into PrP^Sc^ *in vitro* and modulate prion strain properties (40, 41). Moreover, the presence of phosphatidylethanolamine cofactor was necessary to generate infectious forms of recombinant BVPrP (43).

The lack of proteins that bind specifically to BVPrP when expressed in mice is perhaps not surprising, as the structure of the C-terminal globular domain BVPrP^C^ is very similar to that of PrPs from other species and does not contain unique structural motifs that could mediate novel interactions and facilitate prion replication (70). Unlike MoPrP, BVPrP contains a rigid loop in the β2-α2 region, but this microstructure is also present in PrPs from other species such as elk and horses, the latter of which are considered to be quite resistant to prion infection (71, 72). Notably, recombinant BVPrP behaves as a universal substrate for the detection of prion seeds in the real-time quaking-induced conversion (RT-QuIC) assay and spontaneously polymerizes into aggregates much faster than PrPs from other species (26, 32). Therefore, independent of other proteins or cofactors, BVPrP is inherently prone to adopting misfolded conformations. Thus, when paired with our interactome data, the anomalous properties of BVPrP are more likely to be rooted in its intrinsic thermodynamic properties rather than the existence of unique protein-protein interactions.

## Materials and methods

### Mice

Non-transgenic C57BL/6 mice were bred in-house and were originally sourced from Jackson Lab (Stock #000664). B6(Cg)-*Tyr^c-2J^*/J mice (“B6-albino”; Stock #000058) and B6.129S4-*Gt(ROSA)26Sor^tm1(FLP1)Dym^*/RainJ mice (“Flp deleter”; Stock #009086) were also purchased from Jackson Lab. PrP^-/-^ mice on a C57BL/6 co-isogenic background were provided by Adriano Aguzzi (60). Mice were housed in cages of 3-5 animals and were given free access to food and water. The mice were maintained on a 12 h light / 12 h dark cycle. Mice were monitored daily for routine health and checked biweekly for signs of neurological illness. All animal experiments were conducted under an animal use protocol approved by the University Health Network Animal Care Committee (#AUP4263.19).

### Generation of BVPrP knock-in mice

The open reading frame of BVPrP(M109) (GenBank accession numbers AF367624.1 and EF455012.1) was synthesized and then amplified by PCR using the primers 5’-CTATAT<u>GGATCC</u>ACCATGGCGAACCTCAGC-3’ (forward) 5’-CTATAT<u>TCTAGA</u>TCATCCCACGATCAGGAAG-3’ (reverse) for insertion between the *BamH*I and *Xba*I sites of the vector pcDNA3. Generation of the targeting constructs as well as gene targeting of the *Prnp* locus in mouse V6.5 embryonic stem cells using CRISPR/Cas9 technology were performed as described previously (37, 73). Clones with successful gene targeting were expanded and aggregated with diploid CD-1(ICR) mouse embryos at The Centre for Phenogenomics (Toronto, Canada). Chimeric mice were identified by the presence of black patches of fur and were then bred with B6-albino mice to identify those that underwent germline transmission events. The chimeric mice were then bred with Flp deleter mice to remove the selectable marker, and then the offspring of this cross were bred with WT C57BL/6 mice to eliminate the Flp allele. These mice were then intercrossed to generate homozygous knock-in mice. The knock-in mice were maintained by crossing homozygous female with homozygous male mice.

### Tissue harvesting and homogenization

WT C57BL/6 and kiBVM mice at 3-4 months of age were perfused with PBS for 2 min and then brain, spinal cord, heart, lung, muscle, testis, tongue, skin, stomach, and spleen tissue was harvested and snap frozen in liquid nitrogen, followed by storage at −80 °C. Tissues were weighed and then placed in screw cap tubes containing 0.5 mm zirconia beads (BioSpec #11079105Z). Nine volumes of PBS were added to generate 10% (w/v) homogenates. Samples were homogenized three times for 3 min each using a Minilys homogenizer (Bertin Technologies) set at maximum speed, with 5 min incubations on ice in between runs. Some organs such as skin and stomach required additional homogenization to fully break up the tissue. Detergent-extracted brain homogenates were generated by mixing 9 volumes of 10% brain homogenate with 1 volume of 10X detergent buffer [5% (w/v) sodium deoxycholate, 5% (v/v) NP-40 in 1X PBS]. Samples were incubated on ice for 10 min with intermittent vortexing to promote efficient protein extraction and then centrifuged at 5,000x *g* for 5 min at 4 °C. The bicinchoninic acid (BCA) assay (Thermo Fisher #23227) was used to determine total protein concentrations in the detergent-extracted brain homogenates.

### Detergent insolubility assays and thermolysin digestions

To generate detergent-insoluble fractions, detergent-extracted brain homogenates were diluted in 1X detergent buffer [0.5% (w/v) sodium deoxycholate, 0.5% (v/v) NP-40 prepared in DPBS] and then subjected to ultracentrifugation at 100,000x *g* in a Beckman TLA-55 rotor for 1 h at 4 °C. Supernatants were removed and then pellets were resuspended in 1X Bolt LDS sample buffer (Thermo Fisher #B0007) containing 2.5% (v/v) β-mercaptoethanol. The samples were boiled at 95 °C for 10 min and then analyzed by immunoblotting. For thermolysin digestions, 500 μg of detergent-extracted brain homogenate was added to 1X detergent buffer (final volume: 100 µL) containing 50 μg/mL thermolysin (MilliporeSigma #T7902; diluted from a 1 mg/mL stock solution prepared in dH_2_O). This results in a protease:protein ratio of 1:100 in the final reaction. Samples were incubated at 37 °C for 1 h with 600 rpm shaking, and digestions were halted by adding EDTA to a final concentration of 5 mM. Sarkosyl was added to a final concentration of 2% (v/v), and then samples were ultracentrifuged at 100,000x *g* for 1 h at 4 °C. Supernatants were gently removed and then pellets were resuspended in 1X Bolt LDS sample buffer containing 2.5% (v/v) β-mercaptoethanol, boiled, and analyzed by immunoblotting.

### De-glycosylation of proteins using PNGase F

Detergent-extracted brain homogenates containing 100 µg of protein were incubated at 95 °C for 10 min following addition of one volume of 10X glycoprotein denaturing buffer. The samples were cooled on ice, and then 5 μL of 10% NP-40 and 10X GlycoBuffer 2 as well as 1 μL of PNGase F (New England Biolabs #P0704S) were added to make a final reaction volume of 50 μL. Following overnight incubation at 37 °C, reactions were stopped by adding LDS sample buffer (1X final concentration) and boiling the samples at 95 °C for 10 min. De-glycosylated samples were analyzed by immunoblotting.

### Immunoblotting

Samples were run on 10% Bolt Bis-Tris gels (Thermo Fisher #NW00100BOX or NW00102BOX) at 165 V for 35 min. Gels were transferred onto a 0.45 mm Immobilon-P PVDF membranes (MilliporeSigma #IPVH00010) using Tris-Glycine transfer buffer (100 mM Tris-HCl pH 8, 137 mM glycine) at 25 V for 60 min. Following transfer, membranes were blocked for 1 h in blocking buffer [5% (w/v) skim milk in Tris-buffered saline (TBS) containing 0.05% (v/v) Tween-20 (TBST)]. The blocked membranes were then incubated with primary antibody in TBST overnight at 4 °C. The following primary anti-PrP antibodies were used: HuM-D18 (1:5,000 dilution) (74), HuM-D13 (1:10,000 dilution) (74), POM1 (MilliporeSigma #MABN2285; 1:5,000 dilution) (75), HuM-R1 (1:10,000 dilution) (74), SAF-32 (Cayman Chemical #189720; 1:5,000 dilution), and Sha31 (Cayman Chemical #11866; 1:5,000 dilution). The HuM-D18 antibody was produced in-house whereas the HuM-D13 and HuM-R1 antibodies were provided by Stanley Prusiner (University of California San Francisco). The membranes were then washed 3 times with TBST for 10 min each, and then incubated with the appropriate HRP-linked secondary antibody (Bio-Rad #172-1011 or Thermo Fisher Scientific #31414) at a 1:10,000 dilution in blocking buffer for 1 h at 22 °C. Membranes were then washed 3 times with TBST for 10 min and then developed using Western Lightning ECL Pro (Revvity NEL #122001EA) and exposed to x-ray film. For reprobing with an actin antibody, blots were washed with TBST and then treated with 0.05% (w/v) sodium azide diluted in blocking buffer to inactivate the HRP linked to the initial secondary antibody. Blots were then reprobed using the primary anti-actin 20-33 antibody (MilliporeSigma #A5060; 1:10,000 dilution) and an HRP-linked goat anti-rabbit secondary antibody (Bio-Rad #172-1019).

### Time-controlled transcardiac perfusion crosslinking

TcTPC was performed as previously described (53). Briefly, male WT C57BL/6 mice, kiBVM, and PrP^-/-^ mice at approximately 3 months of age were anesthetized with isoflurane and then perfused via the transcardiac route with PBS for 2 min. Following this, mice were perfused with freshly made 2% (w/v) formaldehyde (pH 7.3 in 1X PBS) for 6 min. Successful perfusion was gauged by the development of tail rigidity. The brains were then dissected and post-fixed in 2% formaldehyde for a total of 9 min (includes dissection time). Following fixation, brains were snap frozen in liquid nitrogen and stored at −80 °C.

Immunoprecipitation of crosslinked PrP^C^-containing complexes

KappaSelect beads (Cytiva #17-5458-01) beads were washed, equilibrated, and then conjugated with HuM-D18 Fab by rotation overnight at 4 °C in lysis buffer [150 mM Tris-HCl pH 8.3, 150 mM NaCl, 0.5% (w/v) deoxycholic acid, 0.5% (v/v) NP-40]. The crosslinked brains were homogenized as described above in lysis buffer and then ultracentrifuged at 100,000x *g* for 1 h at 4 °C to remove debris. The brain homogenates were then quantified using the BCA assay and then normalized to the lowest protein concentration amongst all replicates. Homogenates were then incubated with the HuM-D18-conjugated KappaSelect beads overnight at 4 °C in an end-over-end rotator. Following capture, beads were washed 3 times with lysis buffer and then 3 times with a more stringent lysis buffer containing a higher salt concentration (500 mM NaCl). Prior to elution, beads were washed briefly with a buffer containing 10 mM HEPES, pH 8. Captured proteins were eluted by pH drop using a buffer containing 0.2% (v/v) trifluoroacetic acid in 20% (v/v) acetonitrile.

### Sample reduction, alkylation, trypsinization, and iTRAQ labeling

Eluates from the immunoprecipitation of PrP^C^-containing complexes were dried to a volume of 5 µL using a SpeedVac centrifuge set at 37 °C. The dried eluates were denatured by addition of 9 M deionized urea and then reduced by incubation with Tris(2-carboxyethyl)phosphine (TCEP) in 500 mM triethyl ammonium bicarbonate buffer at 60 °C for 30 min. After allowing the samples to cool, sulfhydryl groups were alkylated by treatment with 4-vinylpyridine for 1 h at 22 °C. Urea concentration was diluted to 1.25 M in 500 mM triethylammonium bicarbonate buffer prior to the addition of mass spectrometry grade trypsin (Thermo Fisher Scientific #90057). Trypsin digestions were done overnight at 37 °C. The digested mixture was purified with reverse phase resin, paired with or without a strong cation exchange. A portion of the trypsinized samples were reacted with 8-plex iTRAQ reagents (SCIEX #4390811) according to the manufacturer’s instructions and then combined. Prior to LC-MS/MS, peptides were purified using OMIX C18 pipette tips (Agilent #A57003100).

### LC-MS/MS analysis

Samples were analyzed using an EASY-nLC 1000-Orbitrap Fusion Tribid mass spectrometry platform (Thermo Fisher Scientific) with a 4-hour reversed-phase acetonitrile/water gradient at a flow rate of 300 nL/min. The analytical C18 column (Acclaim PepMap RSLC 100) was 25 cm long with a 75 µm inner diameter. Each LC-MS/MS run consisted of an orbitrap precursor ion scan as well as ion trap (MS2) and orbitrap (MS3) product ion scans within a 3-second window following collision induced dissociation and higher energy collisional dissociation, respectively. For the orbitrap scans, the resolution was set at 60,000.

### Protein identification and quantification

The MS2 data sets were analyzed and converted into peptide sequences with Proteome Discoverer software (version 1.4) using the built-in Mascot and Sequest HT search algorithms and search parameters as previously described (57). The mouse Uniprot database was used as a reference with the manual inclusion of the BVPrP sequence. To stringently filter the mass spectra, the Percolator algorithm within Proteome Discoverer was used to estimate the false discovery rate based on the q-value. The Reporter Ions Quantifier algorithm in Proteome Discoverer was utilized to determine the relative quantification of iTRAQ-labeled peptides using the MS3 data. For the unlabeled datasets, the number of PSMs for the 3 biological replicates of the kiBVM samples, the 3 biological replicates of the WT C57BL/6 samples, and the two biological replicates of the PrP^-/-^ samples were averaged for each identified protein. Hits were then sorted by the average number of PSMs in the kiBVM or WT samples minus the average number of PSMs in the PrP^-/-^ samples. For a protein to be included in the list of identified interactors, its identification had to be supported by at least 2 unique peptides as well as an average kiBVM:average PrP^-/-^ or an average WT:average PrP^-/-^ PSM ratio of at least 3.

### Statistical analysis

Statistical comparisons were conducted using GraphPad Prism software (version 10). iTRAQ reporter ion enrichment ratios were compared using two-tailed unpaired t-tests with a significance threshold of P < 0.05.

## Acknowledgements

The authors thank Rosemary Ahrens for assistance with the mouse perfusions and Stanley Prusiner (University of California San Francisco) for providing the HuM-D13 and HuM-R1 antibodies. Experimental schematics were created using BioRender.com.

## Declarations

### Availability of data and material

Upon publication, the proteomics data will be deposited to the ProteomeXchange Consortium via the PRIDE partner repository. All other data generated or analyzed during this study are included in this published article.

## Author contributions

HA, WSJ, GSU, and JCW contributed to the study conception and design. Material preparation, data collection and analysis were performed by HA, SE, SM, DW, LK, and ES. The first draft of the manuscript was written by HA and JCW, and all authors commented on previous versions of the manuscript. All authors read and approved the final manuscript.

