## Supplemental information for "The Brain Interactome of a Permissive Prion Replication Substrate"

*Running title*: The interactome of bank vole PrP

*Keywords*: prion, protein-protein interactions, mass spectrometry, knock-in mice, bank voles

| | | | | MoPrP interactome | | | | | | | | BVPrP interactome | |
|---|---|---|---|---|---|---|---|---|---|---|---|---|---|
| Rank in MoPrP interactome | Accession | Description (curated) | Gene | # PSM WT (1) | # PSM WT (2) | # PSM WT (3) | # PSM KO (1) | # PSM KO (2) | Avg PSM WT | Avg PSM KO | Avg PSM WT minus Avg PSM KO | Avg PSM kiBVM minus Avg PSM KO | Rank in BVPrP interactome |
| 1 | P04925 | Major prion protein | Prnp | 291 | 105 | 152 | | | 182.67 | 0.00 | 182.67 | 208.00 | 1 |
| 2 | Q6PIC6 | Sodium/potassium-transporting ATPase subunit alpha-3 | Atp1a3 | 247 | 128 | 171 | 50 | 51 | 182.00 | 50.50 | 131.50 | 188.50 | 2 |
| 3 | E9QB01 | Neural cell adhesion molecule 1 | Ncam1 | 175 | 44 | 67 | | 6 | 95.33 | 3.00 | 92.33 | 135.33 | 3 |
| 4 | Q8VDN2 | Sodium/potassium-transporting ATPase subunit alpha-1 | Atp1a1 | 167 | 79 | 113 | 44 | 32 | 119.67 | 38.00 | 81.67 | 119.33 | 4 |
| 5 | Q6PIE5 | Sodium/potassium-transporting ATPase subunit alpha-2 | Atp1a2 | 132 | 92 | 114 | 37 | 35 | 112.67 | 36.00 | 76.67 | 117.00 | 5 |
| 6 | P12960 | Contactin-1 | Cntn1 | 105 | 25 | 35 | 6 | 8 | 55.00 | 7.00 | 48.00 | 71.33 | 6 |
| 7 | Q8BG39 | Synaptic vesicle glycoprotein 2B | Sv2b | 78 | 26 | 30 | 8 | 3 | 44.67 | 5.50 | 39.17 | 50.50 | 7 |
| 8 | A2AFG8 | Neural cell adhesion molecule L1 | L1cam | 59 | 21 | 35 | 7 | 12 | 38.33 | 9.50 | 28.83 | 39.17 | 9 |
| 9 | P60879 | Synaptosomal-associated protein 25 | Snap25 | 50 | 13 | 18 | | | 27.00 | 0.00 | 27.00 | 24.33 | 19 |
| 10 | P46096 | Synaptotagmin-1 | Syt1 | 32 | 17 | 19 | | | 22.67 | 0.00 | 22.67 | 30.00 | 16 |
| 11 | O35136 | Neural cell adhesion molecule 2 | Ncam2 | 49 | 7 | 21 | 6 | | 25.67 | 3.00 | 22.67 | 30.67 | 14 |
| 12 | P14094 | Sodium/potassium-transporting ATPase subunit beta-1 | Atp1b1 | 47 | 18 | 22 | 5 | 8 | 29.00 | 6.50 | 22.50 | 41.83 | 8 |
| 13 | Q5U4C2 | Dipeptidyl aminopeptidase-like protein 6 | Dpp6 | 39 | 9 | 16 | | | 21.33 | 0.00 | 21.33 | 33.00 | 11 |
| 14 | P61264 | Syntaxin-1B | Stx1b | 34 | 17 | 18 | 6 | | 23.00 | 3.00 | 20.00 | 30.33 | 15 |
| 15 | P63101 | 14-3-3 protein zeta/delta | Ywhaz | 37 | 22 | 21 | 10 | 5 | 26.67 | 7.50 | 19.17 | 35.50 | 10 |
| 16 | P62874 | Guanine nucleotide-binding protein G(I)/G(S)/G(T) subunit beta-1 | Gnb1 | 38 | 14 | 20 | 6 | 5 | 24.00 | 5.50 | 18.50 | 27.83 | 17 |
| 17 | E9QKR0 | Guanine nucleotide-binding protein G(I)/G(S)/G(T) subunit beta-2 | Gnb2 | 27 | 13 | 11 | | | 17.00 | 0.00 | 17.00 | 22.67 | 22 |
| 18 | Q02053 | Ubiquitin-like modifier-activating enzyme 1 | Uba1 | 27 | 13 | 22 | 5 | 8 | 20.67 | 6.50 | 14.17 | 22.83 | 21 |
| 19 | Q11011 | Puromycin-sensitive aminopeptidase | Npepps | 23 | 14 | 13 | 6 | | 16.67 | 3.00 | 13.67 | 15.33 | 35 |
| 20 | P09103 | Protein disulfide-isomerase | P4hb | 28 | 5 | 6 | | | 13.00 | 0.00 | 13.00 | 22.00 | 23 |
| 21 | Q8BTU6 | Eukaryotic initiation factor 4A-II | Eif4a2 | 7 | 24 | 8 | | | 13.00 | 0.00 | 13.00 | 10.67 | 68 |
| 22 | Q9JIS5 | Synaptic vesicle glycoprotein 2A | Sv2a | 26 | 17 | 15 | 7 | 6 | 19.33 | 6.50 | 12.83 | 21.50 | 25 |
| 23 | Q810U4-2 | Isoform 2 of Neuronal cell adhesion molecule | Nrcam | 34 | 11 | 12 | 8 | 5 | 19.00 | 6.50 | 12.50 | 30.83 | 12 |
| 24 | Q8BLQ9-2 | Isoform 2 of Cell adhesion molecule 2 | Cadm2 | 31 | | 6 | | | 12.33 | 0.00 | 12.33 | 14.67 | 42 |
| 25 | P62259 | 14-3-3 protein epsilon | Ywhae | 21 | 14 | 7 | 4 | | 14.00 | 2.00 | 12.00 | 16.00 | 32 |
| 26 | E9Q3Q6 | CD166 antigen | Alcam | 24 | 11 | 8 | | 5 | 14.33 | 2.50 | 11.83 | 23.50 | 20 |
| 27 | P28663 | Beta-soluble NSF attachment protein | Napb | 18 | 10 | 13 | | 4 | 13.67 | 2.00 | 11.67 | 14.00 | 46 |
| 28 | P56564 | Excitatory amino acid transporter 1 | Slc1a3 | 19 | 11 | 18 | 4 | 5 | 16.00 | 4.50 | 11.50 | 21.50 | 24 |
| 29 | O35526 | Syntaxin-1A | Stx1a | 13 | 9 | 12 | | | 11.33 | 0.00 | 11.33 | 19.67 | 27 |
| 30 | P46097 | Synaptotagmin-2 | Syt2 | 15 | 9 | 10 | | | 11.33 | 0.00 | 11.33 | 17.00 | 29 |

**Figure S1. The brain interactome of mouse PrP.** Top 30 identified MoPrP-interacting proteins ranked by the average number of PSMs observed in WT C57BL/6 mice minus the average number of PSMs observed in $PrP^{-/-}$ mice (KO). For each protein identified, the average number of PSMs observed in kiBVM mice minus the average number of PSMs from $PrP^{-/-}$ mice as well as the rank in the BVPrP interactome are also shown. Blue-shaded numbers reflect the number of PSMs detected and green-shaded numbers reflect the relative enrichment in WT or kiBVM versus $PrP^{-/-}$ samples.

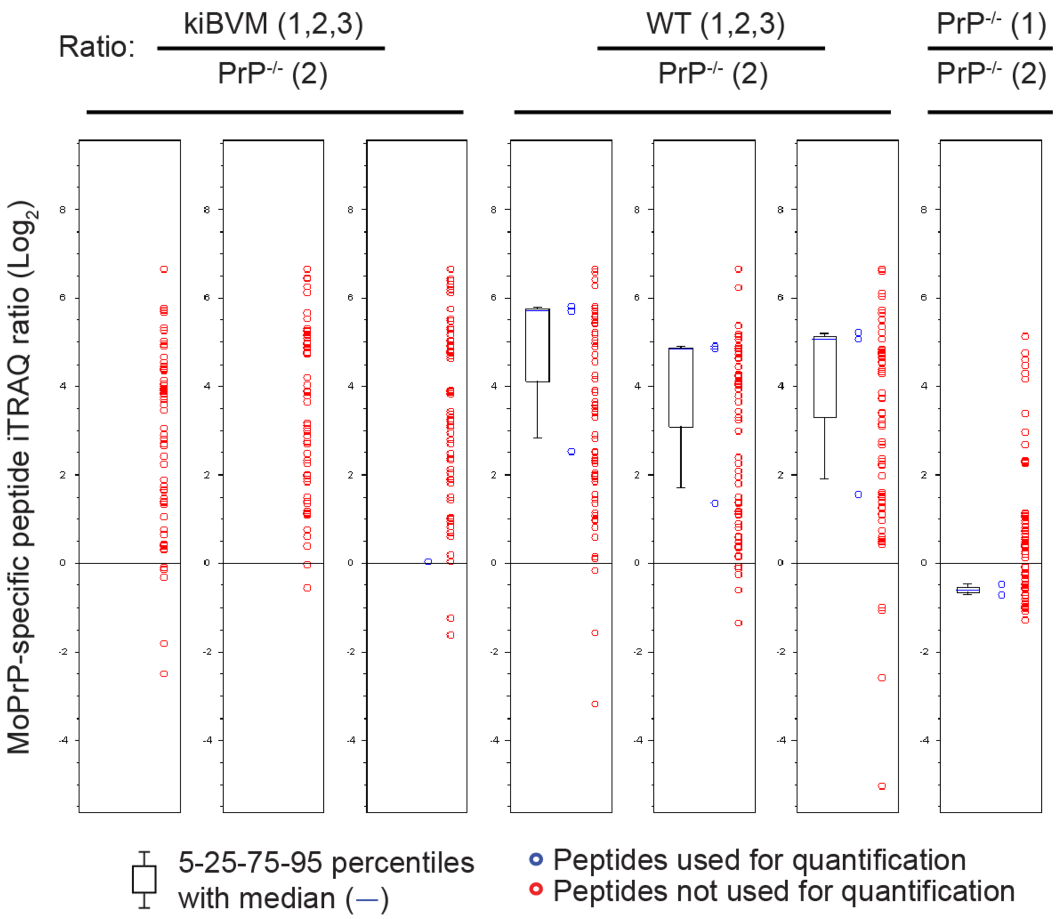


**Figure S2. Analysis of mouse PrP-specific peptides in the iTRAQ-labeled dataset**. Boxplots depicting iTRAQ reporter ion enrichment ratios ($Log_2$) for PrP peptides containing at least one MoPrP-specific amino acid in immunoprecipitated samples from the brains of kiBVM and WT C57BL/6 mice (n = 3 each) as well as the brain of a $PrP^{-/-}$ mouse. For each sample, the enrichment ratio was calculated in comparison to the second $PrP^{-/-}$ sample.

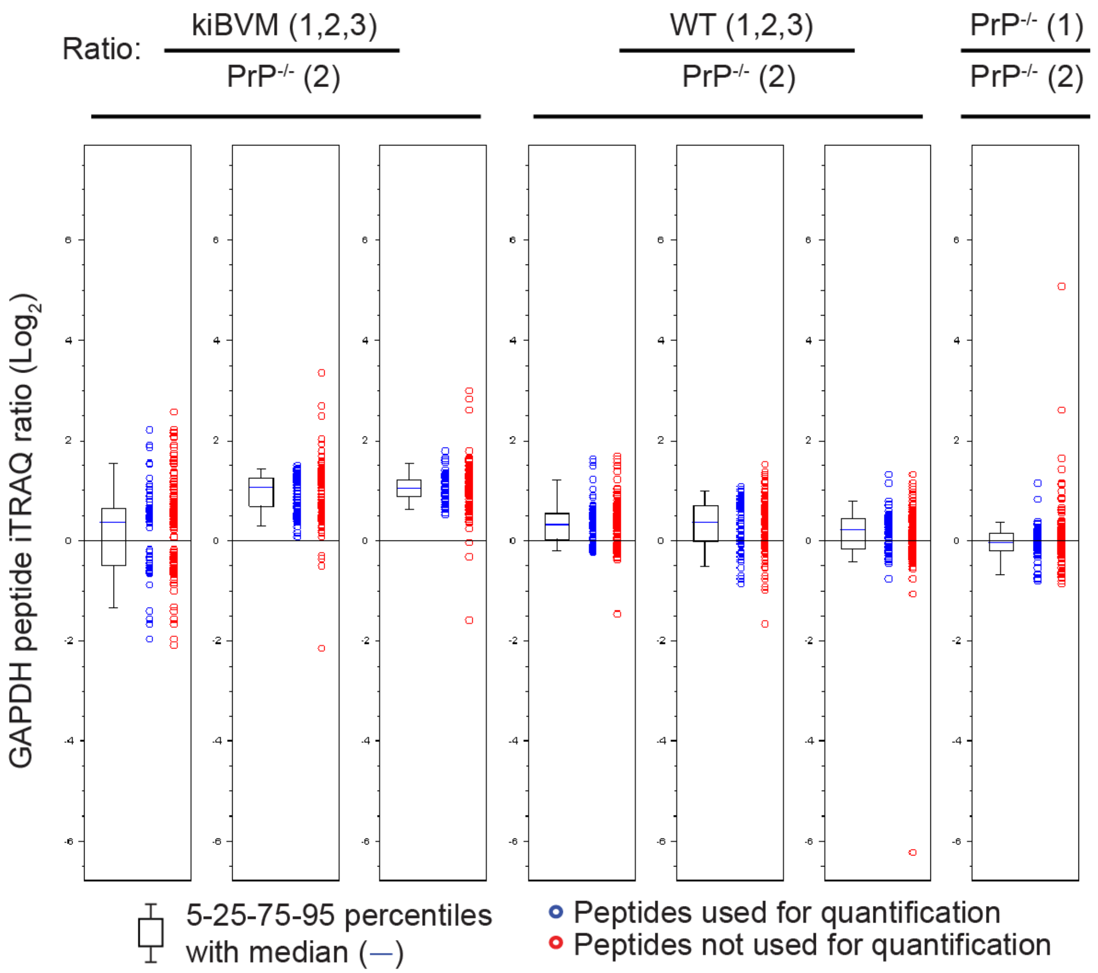


**Figure S3. Non-specific presence of glyceraldehyde-3-phosphate dehydrogenase (GAPDH) peptides in the iTRAQ-labeled dataset**. Boxplots depicting iTRAQ reporter ion enrichment ratios ($Log_2$) for GAPDH peptides in immunoprecipitated samples from the brains of kiBVM and WT C57BL/6 mice (n = 3 each) as well as the brain of a $PrP^{-/-}$ mouse. For each sample, the enrichment ratio was calculated in comparison to the second $PrP^{-/-}$ sample.
